# Persistent Adaptation through Dual-Timescale Regulation of Ion Channel Properties

**DOI:** 10.1101/2025.10.22.684058

**Authors:** Yugarshi Mondal, Ronald L. Calabrese, Eve Marder

**Affiliations:** Volen Center and Biology Department, Brandeis University, Waltham, MA 02454; Biology Department, Emory University, Atlanta, GA, 30322

**Keywords:** intrinsic excitability, homeostatic plasticity, activity-dependent regulation, computational model, high potassium, channel activation curves

## Abstract

Neurons are terminally differentiated cells that adapt to maintain stable function over years, despite encountering a wide range of environmental perturbations. Some adaptations are transient, fading once the perturbation ends. Others are persistent, continuing to influence a neuron’s responses to future challenges even after baseline conditions are restored. These persistent adaptations are especially intriguing because some remain undetectable under normal conditions—only becoming apparent upon re-exposure to a perturbation. Among the many mechanisms that may contribute to persistent adaptation, we investigate one based on the regulation of intrinsic currents. Using a computational model of activity-dependent homeostasis, we show that slow changes in channel density can encode the influence of past experience and shape future responses while rapid shifts in ion channel voltage-dependence provide immediate compensation during perturbations. Together, these dual processes tune a neuron’s intrinsic excitability, enabling persistent adaptation.

**SIGNIFICANCE STATEMENT:** Some neurons display a remarkable property: after adapting to environmental challenges, they return to baseline activity that looks unchanged, even though underlying modifications were made. These modifications may persist, shaping how the neuron responds to future perturbations. We term this phenomenon *persistent adaptation*. Although the underlying mechanisms of persistent adaptations may be diverse, here we demonstrate one possible route: homeostatic regulation of intrinsic excitability. Slow changes in channel density can store a trace of past perturbations, while rapid shifts in channel voltage-dependence can enable immediate recovery during re-exposure.

## INTRODUCTION

Over their lifetimes—often spanning years—neurons encounter countless changes in their external environments that push them away from their typical activity patterns. To maintain stable function, they can, amongst other things, alter their intrinsic excitability. These adaptations may be transient, disappearing once the perturbation ends, or persistent, continuing to shape the neuron’s behavior even after it returns to baseline conditions.

Persistent adaptation is particularly striking because it can remain invisible or cryptic under normal conditions. For example, neurons exposed to repeated high extracellular potassium perturbations recover their bursting rhythm more quickly on subsequent exposures (1, 2), even though their baseline activity appears unchanged. A similar phenomenon occurs during rapid cold hardening (RCH) in insects, where a brief drop in temperature enhances tolerance to future cold stress. Although neural activity appears normal at baseline temperatures, RCH can delay cold-induced spreading depolarization on re-exposure—suggesting that persistent adaptation alters neuronal excitability in ways that are only unmasked by subsequent low temperature perturbations (3, 4).

In both cases, there is indirect evidence that persistent adaptation may involve changes to intrinsic membrane properties. Neurons that resume firing in elevated extracellular potassium exhibit altered responses to current injection (2, 5), while RCH increases in situ Na^+^/K^+^-ATPase activity (3). Because Na^+^/K^+^-ATPase moves net charge across the membrane (outward current), its upregulation can directly affect the neuron’s excitability. However, in both examples, these changes have not yet been mechanistically linked to specific alterations in the ionic currents that set a neuron’s firing properties, leaving open the question of which physiological features support this form of persistent adaptation.

Neurons can regulate their intrinsic currents by modifying two key properties of their ion channels: the number (or density) of channels expressed, and the voltage- and time-dependence of channel gating. Theoretical work has implicated both in adaptation to perturbations. On longer timescales, neurons may adjust channel densities to re-establish stable activity (6-13). On shorter timescales, neurons can adjust their excitability by regulating their channel voltage-dependence (14-25). Such changes can facilitate adaptation to perturbations, and in some circumstances modifications of only a single or pair of currents may be sufficient—for example, during adaptation to high potassium (26). Both mechanisms likely operate in combination to support adaptation, whether transient or persistent.

To investigate how persistent adaptation might arise from the combined influence of voltage-dependence and channel density, we used an activity-dependent homeostatic model in which a neuron adjusts both properties in response to deviations from a target activity level (27). We assumed that changes in channel density occur on slower timescales than changes in voltage-dependence, consistent with the general timescale differences between protein synthesis/degradation and post-translational modifications (27). In this paper we now show that persistent adaptation can emerge from the interaction of these fast and slow processes. Slower changes in channel density can encode a latent trace of past activity as rapid shifts in voltage-dependence provide immediate compensation during perturbations—modeled here as elevated extracellular potassium. This trace is not apparent under baseline conditions but shapes the neuron’s response during subsequent perturbations. Together, these mechanisms can enable persistent adaptation through the coordinated regulation of intrinsic neuronal properties.

## RESULTS

To explore how persistent adaptation can emerge from intrinsic plasticity, we used a conductance-based neuron model with activity-dependent regulation of both ion channel density and voltage-dependence (27).

Briefly, the model consists of a single-compartment Hodgkin-Huxley type neuron with seven intrinsic currents: a hyperpolarization-activated inward current, two calcium currents, one sodium current, three potassium currents, and a leak current. Intracellular calcium fluctuates in response to neuronal activity. Three distinct filters are applied, each designed to extract a different component of the calcium signal: one isolates calcium entry associated with spiking, another captures calcium driven by the underlying slow wave, and a third reflects the overall average intracellular calcium concentration. The decomposition of the calcium signal across distinct timescales reflects a strategy that has been used to analyze intrinsic neuronal dynamics in other biophysical contexts (28). Time-averaged values of each filtered signal are compared to predefined setpoints. These setpoints—specified a priori—represent the neuron’s activity targets (7, 27, 29). When neuronal activity causes sustained deviations from these targets, the model’s activity dependent-regulation mechanism adjusts each intrinsic current’s maximal conductance (reflecting ion channel density) and its voltage-dependent activation and, where applicable, inactivation curves, to bring activity back in line with the targets.

Crucially, the activity-dependent regulation mechanism employed here operates on two timescales: a slow process adjusts maximal conductances (*τ*_*g*_ = 600 s), reflecting protein synthesis and degradation; a fast process shifts the voltage-dependence of activation and inactivation (*τ*_*g*_ = s), mimicking post-translational modifications. All details of the model are provided in Mondal, Calabrese and Marder (27), motivated by the experimental work in Rue, Alonso and Marder (1) and Lee and Marder (2).

In what follows, we take 20 degenerate spontaneously bursting neurons from Mondal, Calabrese and Marder (27) and subject them to a sequence of three increases in extracellular potassium concentration.

### Rapid Recovery during Re-exposure Indicates Persistent Adaptation

Figure 1 illustrates how a model neuron’s ion channel densities and voltage-dependences changed in response to repeated increases in extracellular potassium concentration. Each perturbation elevated extracellular potassium to approximately 2.5× the baseline level and lasted 30 minutes, followed by a 30-minute wash period in baseline conditions (see Methods for details).

**Figure 1.**
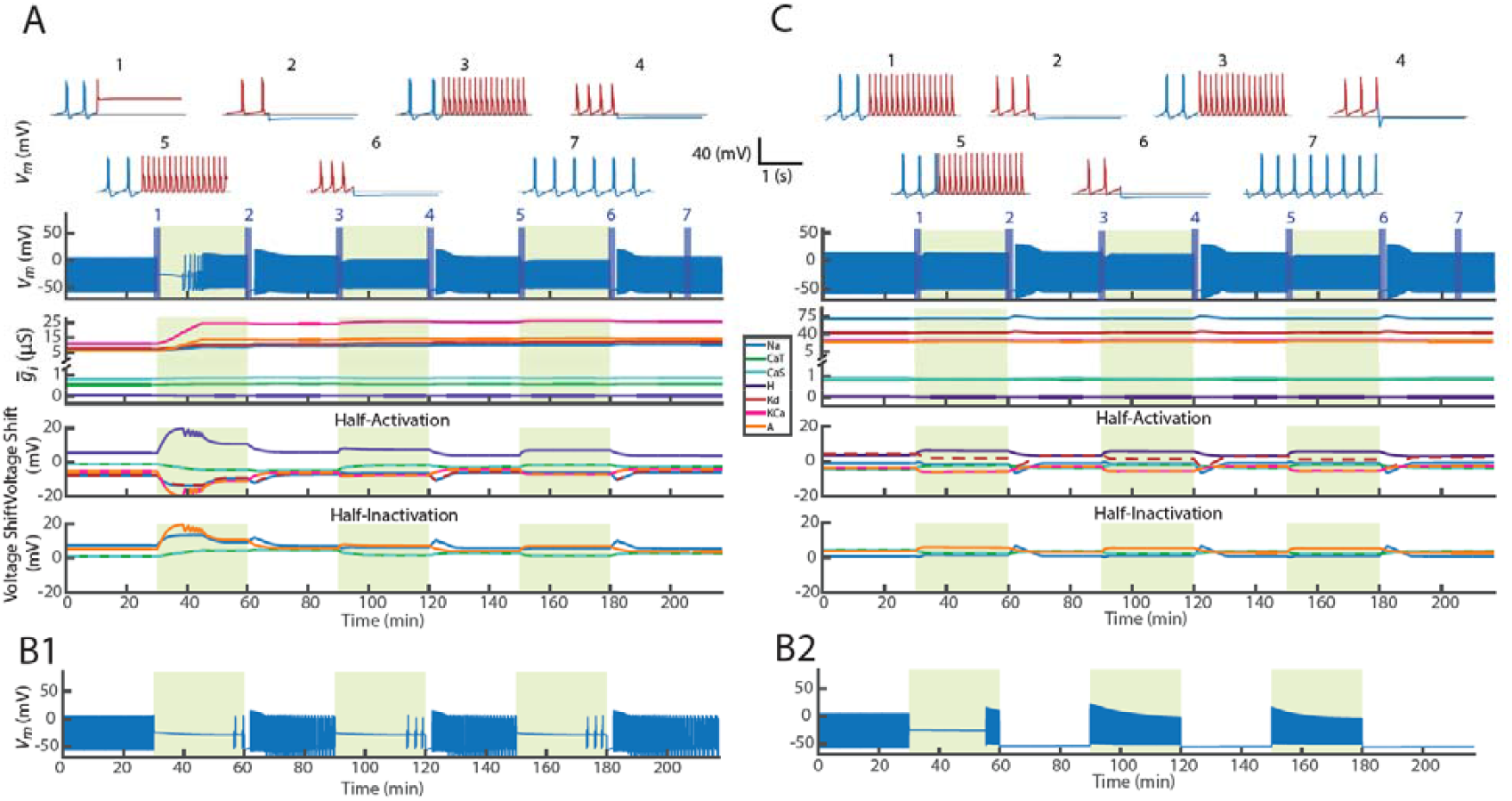
Persistent adaptation emerges from the interaction of slow and fast intrinsic plasticity mechanisms. (**A**) An intrinsically bursting neuron model was subjected to three repeated high extracellular potassium perturbations (green shaded regions). Top two rows show expanded voltage traces (insets 1–7) during selected time points. Red segments indicate activity during perturbation; blue segments indicate activity during wash. Third row shows full voltage trace over time. Fourth row shows changes in maximal conductances for each intrinsic current. Fifth and sixth rows show shifts in voltage-dependent half-activation and half-inactivation voltages. (**B1**) Same model with only voltage-dependence half-(in)activations allowed to vary. (**B2**) Same model with only maximal conductances allowed to change. (**C**) Neuron model with intrinsic parameters pre-adapted to high-potassium perturbation. The plot labels are described as in **A**. Observe that this model exhibited no latency to first burst during the first perturbation.

Before the first perturbation, the neuron fired bursts under baseline conditions (Figure 1-A, inset 1, blue). Upon exposure to high potassium, it entered depolarization block (inset 1, red), but recovered bursting during the perturbation (inset 2, red). Remarkably, when the same perturbation was repeated, the neuron resumed bursting almost immediately (insets 3 and 5, red), indicating that the adaptation established during the first perturbation persisted—constituting a form of persistent adaptation.

Across all models, bursting returned during the wash periods (insets 3, 5, 7, blue). Each wash is accompanied by an initial hyperpolarizing undershoot, suggesting it is an inevitable consequence of releasing the high potassium state. During the perturbation, currents are re-balanced to support bursting in a depolarized environment; when conditions return to normal, this balance is transiently inverted, producing hyperpolarization before stable bursting resumes.

The panels below the voltage trace show that this persistent adaptation coincided with large changes in maximal conductances during the first perturbation, followed by stabilization within a narrower range during later cycles. Intrinsic currents’ half-(in)activations continued to shift across perturbations and washes, reflecting ongoing, fast-timescale tuning of excitability. In this model, both processes were required to reproduce this persistent adaptation. When only half-(in)activations were regulated (Figure 1-B1), the neuron failed to recover during perturbation and resumed bursting only during wash. When only maximal conductances were regulated (Figure 1-B2), the neuron recovered bursting during the first perturbation and exhibited immediate bursting on subsequent ones—suggesting a form of persistent adaptation—but lost bursting stability during wash, instead falling silent. Together, these results suggest that, for this model, fast and slow feedback mechanisms play complementary roles in producing persistent adaptation to repeated potassium perturbations: each alone supports partial recovery, but their interaction enables robust restoration and long-term stability of target activity.

One model in the set of 20 (Figure 1-C) showed no latency to first burst during the first perturbation, suggesting that this model possessed an ion channel configuration already tolerant to the high potassium perturbations unlike the other models that reached this state only after their first exposure.

Figure 2 shows that the pattern of adaptation observed in Figure 1-A was common most models. In 19 out of 20 neurons, the time to first burst during the first perturbation was significantly longer than during subsequent perturbations. The remaining neuron (Figure 2, marked with arrow) exhibited near-immediate bursting even on the first exposure, suggesting it was already well-adapted to the perturbation and that this adaptation persisted on subsequent perturbations (seen in Figure 1-C). In most cases, adaptations were made during the initial perturbation and in all cases, this persisted and enabled faster recovery on subsequent trials.

**Figure 2.**
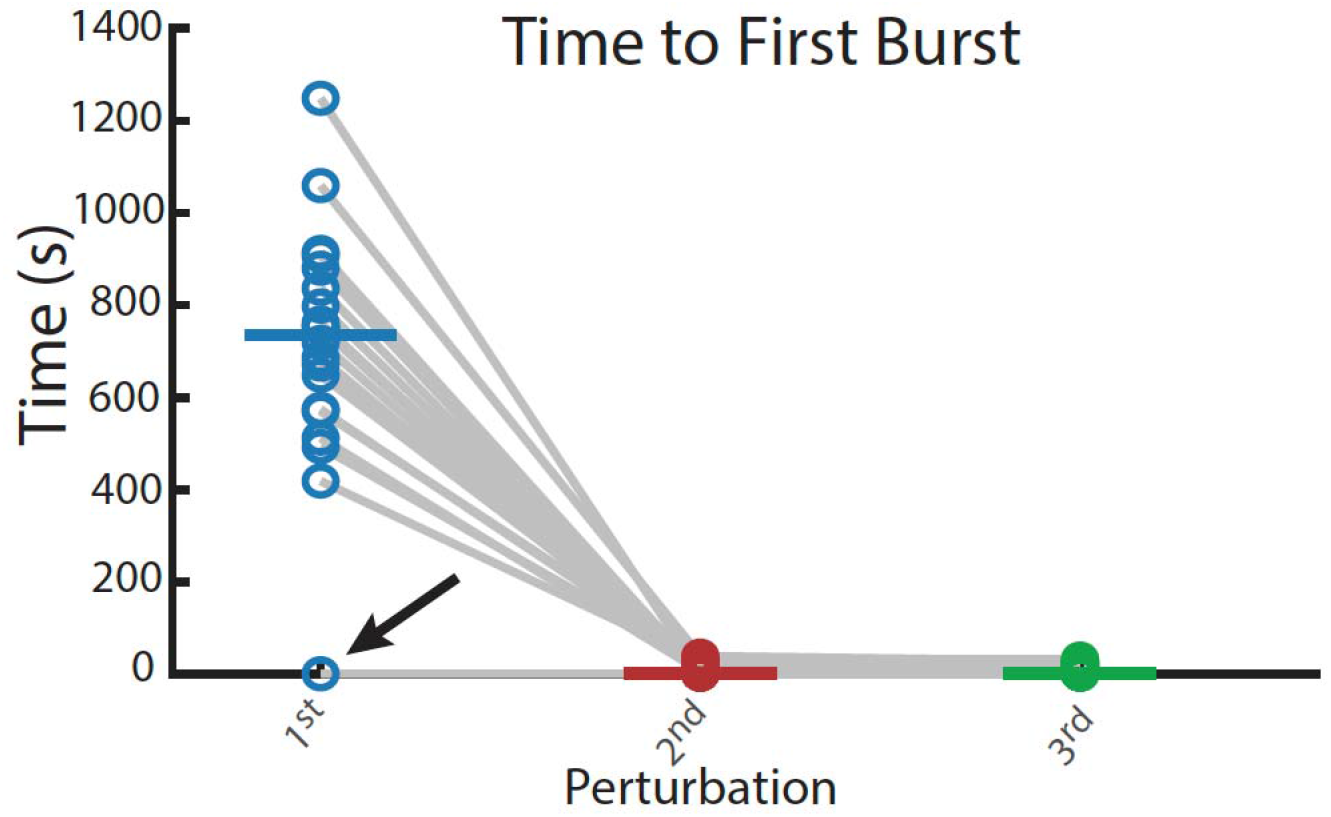
Persistent adaptation reveals itself as reduced time to first burst across repeated perturbations. Each dot represents time to first burst during each of three repeated high-potassium perturbations across 20 model neurons. Each gray line tracks a single model across perturbations. The arrow marks the model that was already adapted to the perturbations (Figure **1-C**).

### First Perturbation Triggers Large Maximal Conductance Shifts That Then Stabilize

In Figure 1-A, we saw that the maximal conductances changed during the first perturbation and stabilized during subsequent perturbation and wash cycles. This, in fact, was representative of most models’ behavior. To capture this, we constructed a simplified geometric representation, schematized in Figure 3-A1. We first collected the neuron’s full set of maximal conductances at seven time points: just before the first perturbation, at the end of the first perturbation, just before the second perturbation, at the end of the second perturbation, and so on—capturing its intrinsic state throughout all perturbation and wash phases up to the end of the third perturbation. We then computed six displacement vectors by subtracting each point from the one that immediately follows it—i.e., computing next minus initial. Each resulting vector quantifies how much—and in what direction—the conductance configuration changed during that time interval.

**Figure 3.**
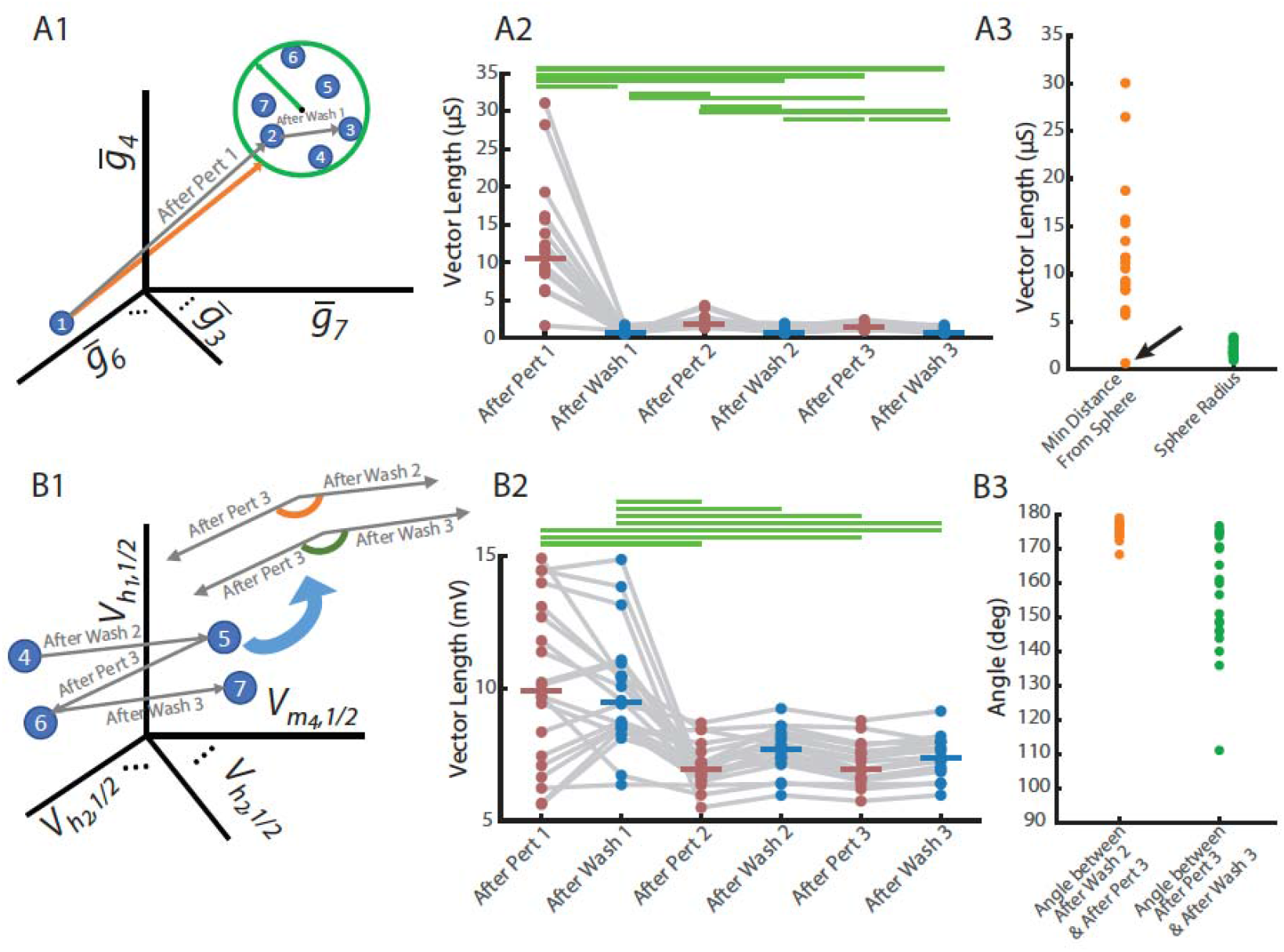
Persistent adaptation involves a large initial conductance shift followed by switching of voltage-dependent properties across repeated perturbations. (**A1**) Schematic of conductance space trajectory for a representative neuron. Each numbered point corresponds to the neuron’s set of maximal conductances at a time point during the perturbation or wash cycles. The first two segments of the trajectory are explicitly shown (gray vectors). The orange vector connects the initial state (point 1) to the closest point on the surface of the minimum bounding sphere (green circle), which is centered at the average of points 2–7. (**A2**) Vector lengths for all 20 models showing the magnitude of change in maximal conductances across successive perturbation and wash cycles. Horizontal bars indicate statistically significant pairwise differences identified using Kruskal–Wallis followed by Dunn’s test with Bonferroni correction (adjusted p < 0.05). (**A3**) Summary measurements in conductance space for each model. Green points show the radius of the final conductance bounding sphere (defined in **A1**), which encloses the conductance states following the first perturbation (points 2–7). Orange points indicate the minimum distance from the initial conductance state (point 1) to the outside of this sphere. The arrow marks the model that was already adapted to the perturbations (Figure 2). (**B1**) Schematic of half-(in)activation space for the same representative model. Each numbered point represents the neuron’s set of half-(in)activations at a specific time point during the perturbation or wash cycles; only the final four points in the sequence are shown here. Labeled vectors indicate the trajectory segments used in the angle analysis. To measure angles, the two vectors in each sequence were positioned tail-to-tail. The angle for the perturbation–wash– perturbation sequence (After Wash 2 to After Perturbation 3) is shown in orange, and the angle for the wash–perturbation–wash sequence (After Perturbation 3 to After Wash 3) is shown in green. (**B2**) Vector lengths for all 20 models showing the magnitude of change in half-(in)activations across successive perturbation and wash cycles. Statistical comparisons were performed using Kruskal–Wallis followed by Dunn’s test with Bonferroni correction; horizontal bars denote significant pairwise differences (adjusted p < 0.05). (**B3**) Angles, as defined in (**B1**), for all 20 model neurons. Each point represents the measured angle for one model in either the perturbation–wash–perturbation sequence (orange) or the wash–perturbation–wash sequence (green).

In Figure 3-A2, these vectors are labeled on the x-axis. For example, “After Pert 1” reflects the change in conductances from just before the start of the first perturbation to just before it ended—containing the full duration of the first high-potassium exposure. “After Wash 1” reflects the change from the end of the first perturbation to just before the second perturbation, and so on through the third perturbation and its wash. In total, these six vector lengths summarize how much the neuron’s conductance profile shifted during each perturbation and wash phase. As shown in Figure 3-A2, the largest changes occurred during the first perturbation, followed by relatively minor adjustments in subsequent phases.

To simplify the patterns observed in Figure 3-A2, we constructed a scheme to represent how conductance states stabilized over time. Specifically, we computed a bounding sphere that enclosed the final 6 points in conductance space—schematically illustrated by points 2 through 7 in the illustration (Figure 3-A1). This sphere represented the region of maximal conductances occupied by the neuron once it had adapted. We then measured the shortest distance from the conductance state immediately preceding the first perturbation to the boundary of this sphere. If this distance exceeded zero, it indicated that the first state lay outside the region occupied in later perturbation-wash cycles (see Methods).

This was true for all models tested, supporting the idea that a major, shift in conductance configuration occurred during the first perturbation, followed by stabilization within a narrower regime (Figure 3-A3). Notably, in 19 out of 20 models, the distance from the pre-perturbation-1 conductance state to the center of the adapted region exceeded the radius of the bounding sphere. This means the neuron’s initial conductance profile lay not just outside the range of its later configurations, but well beyond it. In these cases, this distance was greater than at least twice the radius— indicating that the neuron began far from the set of states it would go on to occupy.

The remaining model corresponded to the preadapted case discussed earlier (Figure 1-C). As noted, firing in this model recovered almost immediately after the first perturbation. Its initial conductance state lay just outside the adapted region (Figure 3-A3, arrow), unlike the other models that began far away. Unlike the others, only a small adjustment was required—underscoring how proximity to the adapted regime can sharply reduce the scale of change needed for recovery.

### Persistent Adaptation Involves Revisiting Half-(In)Activation Profiles

As seen in Figure 1-A, the half-(in)activation parameters did not settle, but instead alternated between different configurations during perturbation and wash phases. In fact, these trajectories suggest a switching behavior—moving toward one configuration of half-(in)activations during perturbation and returning to another during wash. To quantify the switching behavior observed in half-(in)activation parameters, we constructed another geometric representation, schematized in Figure 3-B1. Specifically, we asked whether the neuron departed each regime along a trajectory which was reverse of how it entered. For wash, we focused on the transition from Wash 2 to Perturbation 3 and compared it to the following transition from Perturbation 3 to Wash 3. These two transitions—represented by the vectors labeled After Perturbation 3 and After Wash 3 in Figure 3-B2—capture the last full excursion out of and back into the wash condition. The angle between these vectors were computed by aligning them tail-to-tail in high-dimensional space (Figure 3-B1). Similarly, for perturbation, we compared the transition from Perturbation 2 to Wash 2 with the following transition from Wash 2 to Perturbation 3—represented by the vectors After Wash 2 and After Perturbation 3—and again measured the angle between them.

In all models, these angles exceeded 90 degrees, indicating that the system reversed direction between successive visits to the same regime (Figure 3-B3). When combined with the observation that vector magnitudes remained stable (Figure 3-B2), this suggests that neurons alternated between two half-(in)activation configurations— one for wash, one for perturbation—rather than converging on a single set of parameters.

Taken together, Figure 3 suggests that, following initial adaptation driven by changes in channel expression, the neurons settled into a stable channel repertoire, while half-(in)activations shifted between two configurations to maintain bursting across both wash and perturbation conditions. Figure 1-A shows this happening in a specific neuron model.

### Persistent Adaptation Can Be Stored in Maximal Conductances

The foregoing results suggest that the neuron’s persistent adaptation arises from the changes in expression of ion channels induced by the initial perturbation. To test whether this persistence is stored in the maximal conductance configuration, we re-exposed each model to the same sequence of perturbations, starting from two distinct sets of initial conditions. In the first condition (Figure 4-A), we preserved the final maximal conductances from the original simulation but reset all half-(in)activation voltages to their original, pre-perturbation values. In the second condition (Figure 4-B), we did the opposite: we kept the final half-(in)activation voltages but restored the original maximal conductances. In both cases, the homeostatic mechanism remained active throughout. Independently resetting one parameter set while carrying over the other from the end of the original simulation allowed us to isolate the contributions of maximal conductances versus half-(in)activation voltages in supporting rapid re-adaptation to perturbations.

**Figure 4.**
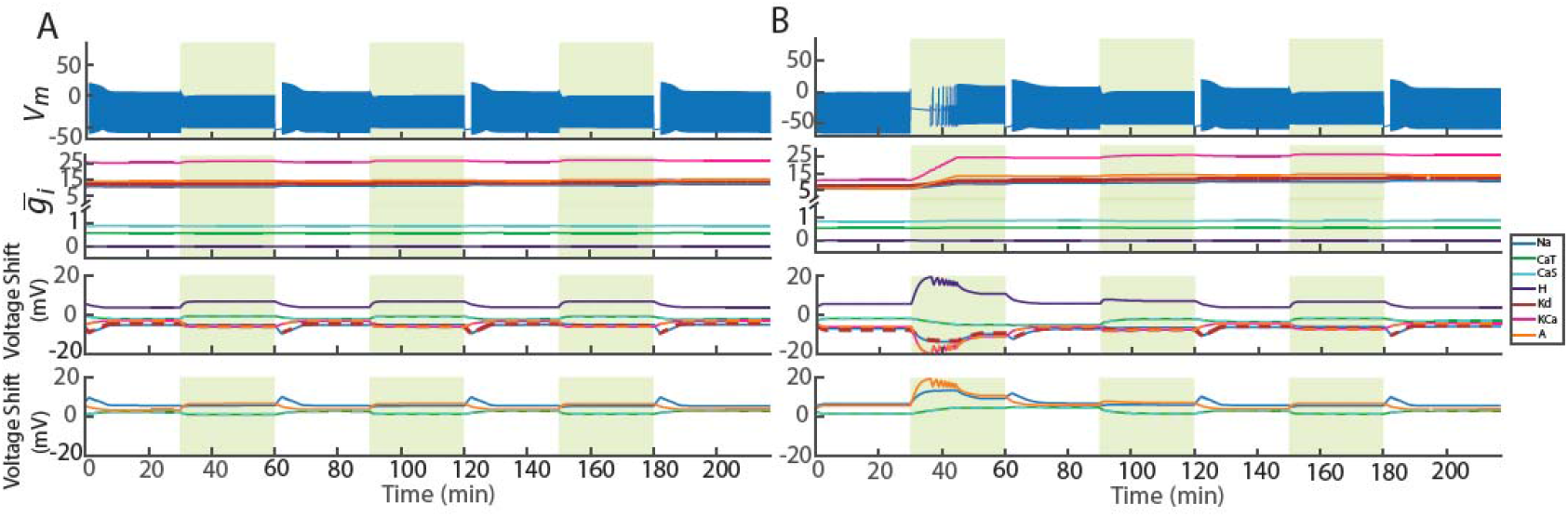
Testing how conductance and voltage parameters store persistent adaptation. (**A**) The model from Figure 1-A was first reconfigured to retain the final maximal conductances from the end of the original simulation while resetting all half-(in)activation voltages to their pre-perturbation values. It was then re-exposed to the same perturbation sequence as in Figure **1-A**. Rows are arranged as in Figure **1-A**. (**B**) The same model was reconfigured to preserve the final half-(in)activation voltages while restoring maximal conductances to their original pre-perturbation values, and then re-exposed to the same perturbation sequence. Rows are arranged as in Figure**1-A**.

The results revealed a striking asymmetry. When the final maximal conductances were retained but the half-(in)activations were reset (Figure 4-A), the model rapidly re-established bursting during the first perturbation—suggesting that the conductance profile alone was sufficient to support immediate adaptation. In contrast, when the final half-(in)activation voltages were retained but the maximal conductances were reset (Figure 4-B), the model exhibited depolarization block during the first perturbation and required time to readapt —resembling the behavior seen in Figure 1-A during initial exposure. This suggested conductance profiles shifts were also perhaps a necessary condition for adaptation.

For all tested models, those initialized with the final maximal conductances (Figure 5, Purple) resumed bursting almost immediately during the first perturbation, as in Figure 4-A. This indicates that the conductance profile alone was sufficient to support immediate adaptation. In contrast, 18 of the 20 models initialized with the final half-(in)activation voltages but restored to their original, pre-perturbation maximal conductances (Figure 5, Green) required substantially more time to recover bursting— mirroring the slow response seen during the first perturbation in the original simulations (Figure 4-B). This pattern suggests that, in most cases, slow changes in maximal conductances were also necessary for immediate adaptation. Of the two exceptions (Figure 4-A, arrow), one was the preadapted case (Figure 1-C), where the original conductances were already tuned for the perturbation. The remaining adapted rapidly despite the conductance reset, implying that in some cases the original conductance repertoire can still support immediate adaptation if paired with a different half-(in)activation configuration.

**Figure 5.**
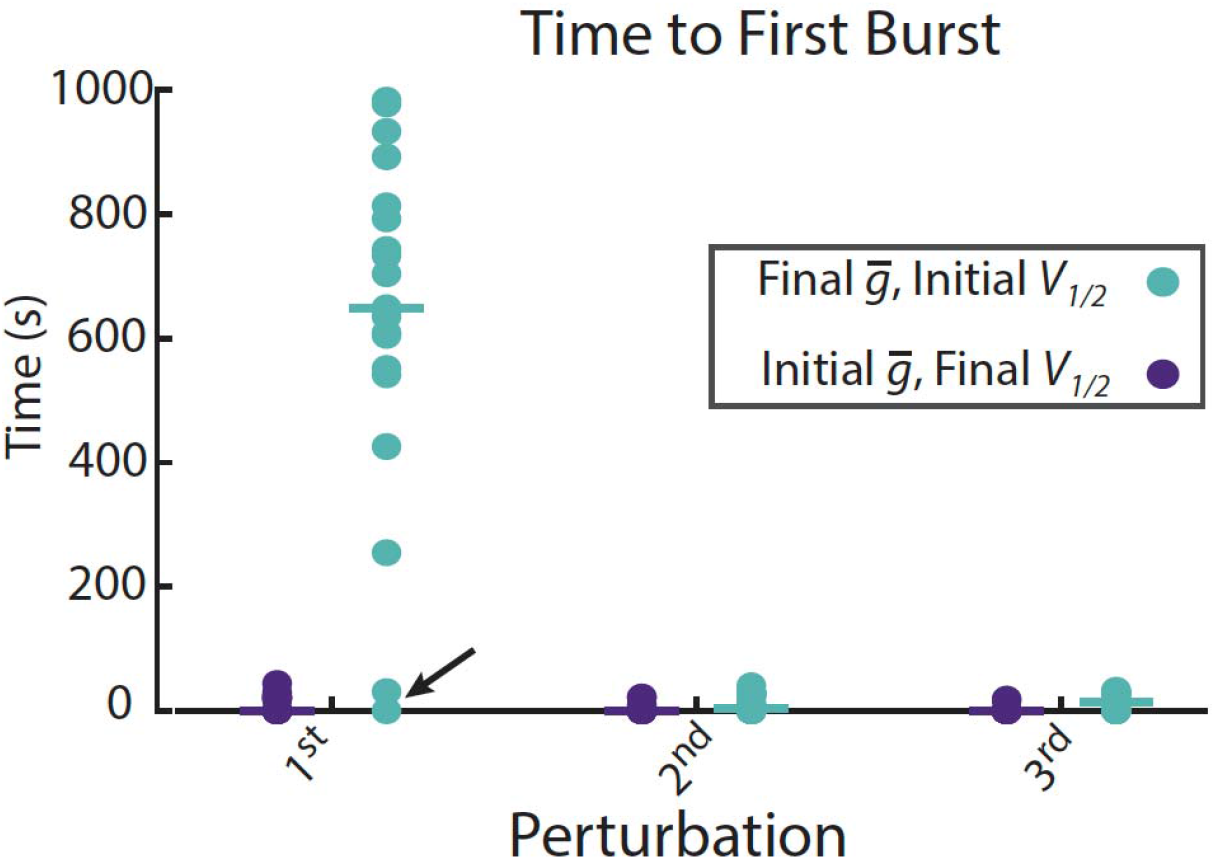
Time to first burst across repeated perturbations when maximal conductances and half-(in)activations are reconfigured. This analysis replicates Figure **2** for the same 20 model neurons, but with parameter rearrangements described in Figure **4**. Models were reconfigured either to retain final maximal conductances while resetting half-(in)activation voltages to their pre-perturbation values (purple), or to retain final half-(in)activation voltages while resetting maximal conductances (cyan) and were then re-exposed to the same sequence of perturbations. The arrow marks the models that were already adapted to the perturbations.

To further interpret the differences in adaptation times, we revisited the geometric framework from Figure 3-A1 to analyze how maximal conductance configurations evolved during re-exposure. Using the same approach to define each model’s stable regime in conductance space, we constructed a bounding sphere around the conductance states observed after the initial perturbation and measured how close the conductance state at the beginning of re-exposure lay to this region. As shown in Figure 6-A3, 20 of 20 models initialized with their final maximal conductances (purple) began near the trajectory of subsequent states during re-exposure. In contrast, 17 of 20 models reset to their original pre-adaptation maximal conductances (cyan) often started far outside this region and had to traverse large distances in conductance space to converge toward it. The remaining three models (Figure 6-A3, arrow) began re-exposure already close to the preadapted state. Two of these corresponded to the rapid-adapting cases described above. The third case did achieve adaptation during the first perturbation, but the conductance parameters did not stabilize into a confined region of conductance space until the second perturbation.

**Figure 6.**
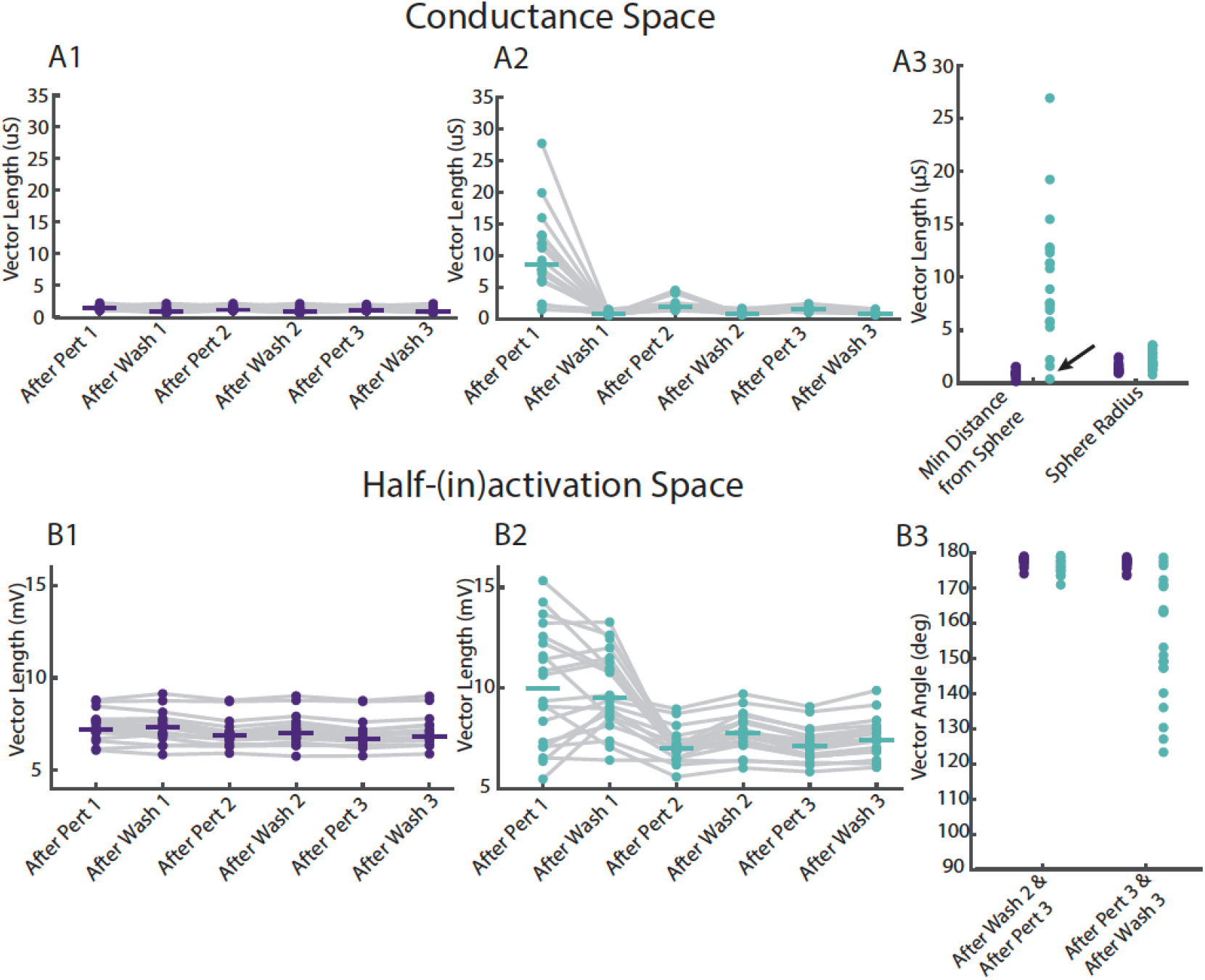
Trajectories in parameter space during re-adaptation of reconfigured models. The neuron models were reconfigured in the following ways: either initialized with final maximal conductances and pre-perturbation half-(in)activation voltages (purple) or initialized with pre-perturbation maximal conductances and final half-(in)activation voltages (cyan). Trajectories of the reconfigured models from Figure **4** during re-exposure to the same sequence of high-potassium perturbations. (**A1**) Vector lengths in maximal conductance space after each perturbation and wash phase for models initialized with final maximal conductances and pre-perturbation half-(in)activation voltages (same analysis as in Figure **3-A2**). (**A2**) Vector lengths in maximal conductance space after each perturbation and wash phase for models initialized with final half-(in)activation voltages and pre-perturbation maximal conductances (same analysis as in Figure **3-A2**). (**A3**) A comparison of reconfigured models’ vector distances in maximal conductance space at the start of re-exposure, showing each model’s minimum distance to the bounding sphere defined by post-adaptation states (left) and the radius of that sphere (right) (same analysis as in Figure **3-A3**). The arrow mark models whose pre-perturbation states lay close to the post-adaptation sphere. (1) Vector lengths in half-(in)activation voltage space after each perturbation and wash phase for models initialized with final maximal conductances and pre-perturbation half-(in)activation voltages (same analysis as in Figure **3-B2**). (**B2**) Vector lengths in half-(in)activation voltage space after each perturbation and wash phase for models initialized with final half-(in)activation voltages and pre-perturbation maximal conductances (same analysis as in Figure **3-B2**). (**B3**) A comparison of reconfigured models’ angles between displacement vectors of half-(in)activation parameters during transitions between perturbation and wash phases (same analysis as in Figure **3-B3**).

We also examined the dynamics of half-(in)activation voltages during re-exposure by measuring the angles between successive displacement vectors. For each model and initialization, the lengths of the displacement vectors during the final transitions into and out of the wash and perturbation phases were consistent, allowing us to focus on direction rather than magnitude, as in Figure 3-B1 (Figure 6-B1 and 6-B2). Across both re-initialization conditions, these transitions showed directional reversals in half-(in)activation parameter space—though to varying extents. Models initialized with their final maximal conductances (Figure 6-B3, purple) exhibited angles tightly clustered near 180 degrees, indicating consistent reversals in trajectory: the neuron departed from and returned to similar half-(in)activation configurations, whether starting from wash or perturbation phase. In contrast, models reset to their original conductances (Figure 6-B3, cyan) showed greater variability—particularly during transitions into and out of the final wash. This increased dispersion might have been expected though because it mirrors the behavior seen during initial adaptation (Figure 3-B3), when models were still establishing a stable switching strategy. Together, these results suggest that while half-(in)activation parameters enable flexible modulation of excitability across environmental contexts, the underlying persistence of the adaptive response is encoded in the maximal conductance profile.

## DISCUSSION

Our results show that persistent adaptation to repeated perturbations can emerge from a coordinated interplay between slow, expression-based changes in maximal conductances and fast, reversible shifts in channel voltage-dependence. Generally speaking, during an initial high-potassium challenge, model neurons traversed large distances in conductance space, leaving their pre-perturbation profiles to settle into a distinct, stable region. This shift effectively “stored” the adaptive solution, allowing the neuron to begin subsequent perturbations already poised for rapid recovery. Across alternating perturbation and wash phases, voltage-dependent parameters then adjusted dynamically to fine-tune excitability on short timescales.

Within this general framework, analysis of multiple models also uncovered important exceptions that reveal multiple viable routes to adaptation. One model was preadapted, already possessing a conductance repertoire close to the stable regime. Another model was “unlocked” by pairing their original conductances with a different half-(in)activation configuration, enabling immediate adaptation without prior slow changes. Still another model achieved activity adaptation before conductance stability, converging only after an additional perturbation. These cases show that while slow conductance changes were sufficient, they weren’t always necessary.

### The Importance of Dual-Timescale Control

The complementary division of labor—slow conductance reconfiguration giving rise to long-lived stability and fast voltage-dependence shifts allowing real-time tuning— suggests a mechanism in which persistent adaptation is not simply a property of one mechanism, but of at least two processes interacting. When only voltage-dependence was regulated (Figure 1-B1), neurons failed to recover during perturbations, regaining bursting only in wash phases. When only maximal conductances were regulated (Figure 1-B2), neurons showed rapid recovery and persistent adaptation across perturbations but lost bursting stability during wash. This demonstrates a broader principle: for robust performance, activity-dependent adaptation should operate across multiple timescales. Rapid mechanisms, such as phosphorylation, can tune excitability within seconds to minutes, but risk destabilizing baseline firing or masking the cues that guide slower, integrative processes. Slow homeostatic regulation stabilizes baseline activity over hours to days but responds too sluggishly to adapt quickly to perturbations. In our model, combining the dual mechanisms suggest a solution to this trade-off: slow, expression-driven changes place the neuron in a region of conductance space where fast voltage-dependence shifts are both effective and stable. The fast mechanism manages transient fluctuations without driving runaway excitability, while the slow mechanism consolidates the adaptive state. The importance of complementary intrinsic homeostatic mechanisms operating at different timescales is consistent with observations in synaptic homeostatic plasticity, where multiple pathways operating on different timescales may potentially coordinate to stabilize synaptic transmission (30, 31).

### Conceptual Assumptions

Our results rest on several conceptual assumptions about how homeostatic regulation might occur. First, the model assumes that homeostasis acts to maintain a fixed activity set point, rather than a broader acceptable range. In biological neurons, a range-based regulation scheme could allow activity to drift within limits, potentially altering the character of the adaptations observed here. Second, the modeled neurons are effectively “naïve,” with no history of prior perturbations before those simulated. Neurons that have already adapted to similar challenges may start closer to an adapted state, limiting the apparent persistent effect. Third, the 20 neurons studied represent a homogeneous subset of the broader degenerate solution space (see Methods). While this controlled selection isolates a clean example of persistent adaptation via timescale separation, it does not imply that all parameter combinations behave similarly. Instead, the findings illustrate one plausible mechanism among many within the solution space. Finally, we assume a simplified representation of excitability where all the firing dynamics are determined by a sets of voltage- and time-dependent ion channel processes. Actually, in biological neurons, excitability arises from a richer interplay of mechanisms —for instance specialized channels that support rapid firing (32) or metabolic feedbacks that couple activity to cellular energy use (33). Exploring these assumptions in future work—such as varying the homeostatic rule to target a range or initializing from partially adapted states—could clarify the generality of the observed dynamics. Additional technical assumptions associated with the model are evaluated in Methods.

### Persistent Adaptation and Cryptic Variability

Persistent adaptation, as observed in the simulations, reflects an important principle: the model neuron can settle into a new configuration that still satisfies its activity target. This contrasts with a simple homeostatic framework, where one might expect a perturbed neuron to eventually return to its original state, leaving no lasting trace once the perturbation ends. Persistent adaptation arises because many different combinations of intrinsic currents create a space of multiple valid solutions that all generate the same firing pattern. When homeostatic regulation navigates this degenerate solution space, neurons that appear similar at baseline can in fact rely on distinct intrinsic current profiles—a phenomenon known as cryptic variability.

Such cryptic variability can have potentially advantageous or maladaptive consequences. On the one hand, the existence of multiple solutions means that adaptation could, in principle, place the neuron in a maladaptive state. For instance, in neuropathic pain, persistent intrinsic modifications mean that even after the acute injury heals, the neurons remain persistently responsive, underpinning chronic pain sensations (34, 35). On the other hand, cryptic variability can be potentially advantageous: in the model, once adaptation has occurred, the fast mechanism allows the neuron to move rapidly among multiple target-satisfying solutions, enabling quicker recovery from repeated perturbations. The qualifier “potentially” is important because an ion channel density state that enhances recovery from one perturbation may, at the same time, increase vulnerability to a different perturbation. These cryptic changes at the individual-cell level may then propagate to the circuit scale, where intrinsic tuning across neurons collectively shapes network excitability and function (36-40). The computational consequences of these cryptic changes would be interesting to follow up on (41, 42).

### Persistent Adaptation as Intrinsic Metaplasticity

Persistent adaptation leaves the neuron in a new intrinsic configuration that changes how it responds to future perturbations. In these simulations, the first perturbation often drives the neuron into depolarization block. Then slow adjustments in the maximal conductances “prime” the neuron, so that during subsequent perturbations, fast half⍰(in)activation shifts immediately fine⍰tune intrinsic currents to recover the target activity. In this way, the neuron’s prior adaptation alters the rules for its future plasticity—embodying plasticity of plasticity, or metaplasticity.

The idea that plasticity itself can be plastic was formalized at the synapse by the Bienenstock–Cooper–Munro (BCM) framework (43). Subsequent work has shown how this might be implemented biophysically through, for instance, synaptic scaling (44) or changes in postsynaptic receptor subunit composition (45). Our results suggest that intrinsic homeostatic plasticity may play a parallel role, providing a biophysical mechanism for metaplasticity through changes in intrinsic excitability, mediated through alterations in channel density and voltage-dependence.

## METHODS

The homeostatic model consisted of a conductance-based neuron, a set of equations that sensed fluctuations in intracellular calcium concentration, and a mechanism that adjusted maximal conductances and channel half-(in)activation voltages in response to deviations from prespecified target calcium levels. These calcium targets served as proxies for the neuron’s target activity pattern. All parameters— including reversal potentials, gating variable time constants, (in)activation curves (with their initial half-activation and half-inactivation voltages), calcium sensor parameters, target calcium levels, and the parameters governing how maximal conductances and half-(in)activations are modulated, as well as the integrator’s parameters—are described in detail in Mondal, Calabrese and Marder (27). The 20 models analyzed in the present study are the same 20 models investigated in that work and all had a burst period within 20% of the average.

Perturbations were administered as in Mondal, Calabrese and Marder (27). Briefly, increasing the extracellular potassium concentration altered both the potassium reversal potential and the leak reversal potential. Using the Nernst equation, the potassium reversal potential was shifted from –80 mV to –55 mV. Using the method described in Mondal, Calabrese and Marder (27), the leak reversal potential was shifted from –50 mV to –32 mV.

### Burst Onset Measurements

The latency from perturbation onset to the start of regular bursting was measured using a custom Python function that leverages the Electrophys Feature Extraction Library (eFEL) distributed with BluePyOpt (46). For each perturbation trial, the voltage trace and corresponding time vector were formatted as an eFEL trace object and passed to the get_feature_values function together with one of eFEL’s predefined extractable features—peak_time in this case—using eFEL’s default spike detection parameters. This returned the times of all detected spike peaks. Inter-spike intervals (ISIs) were then computed, and the onset of regular bursting was defined as the first spike occurring after the last ISI greater than 5 s. If no ISIs exceeded this threshold, the time of the first detected spike was taken as the burst onset. This procedure was applied to each perturbation trial of each neuron to produce the latency values plotted in Figure 2A.

### Quantifying Parameter Space Trajectories

To quantify changes in intrinsic properties over time, we extracted each neuron’s full set of maximal conductances (or half-(in)activation voltages) at seven predefined simulation time points spanning perturbation and wash phases: just before each of perturbations 1, 2, and 3; at the end of each perturbation; and after the final wash (seven points total). Values were averaged over a 20 s window (from 30 s to 10 s before the event) to avoid selecting a single time point arbitrarily. The neuron’s parameters were already stable over this period, so the averaging does not affect the measured values.

For each neuron, trajectories in conductance space and half-(in)activation space were characterized by computing vectors between successive points, obtained by subtracting one point from the next. The Euclidean distance of each vector was calculated by squaring each element, summing across dimensions, and taking the square root. This yielded six vector lengths per neuron for maximal conductances, and another six for half-(in)activations.

To characterize the adapted conductance regime, we analyzed the six states following the first perturbation (points 2–7). Their mean was taken as the center of the adapted region in conductance space. The Euclidean distance from each point to this center was then computed, and the multidimensional sphere radius was defined as the largest of these distances—representing the smallest sphere centered at the mean that enclosed all post-adaptation states. We then measured the Euclidean distance from the conductance state prior to the first perturbation to the center of the adapted region. Subtracting the sphere radius from this distance yielded how far outside the final adapted regime the initial state lay.

To characterize the behavior in half-(in)activation space, we examined whether the trajectory in half-(in)activation space reversed direction between repeated visits to the same condition. We focused on two sequences late in the simulation: the final wash– perturbation–wash cycle, and the final perturbation–wash–perturbation cycle. In each case, we treated the two consecutive transitions as vectors in parameter space and calculated the angle between them using the inverse of the dot-product relationship between vector direction and angle. Angles were reported in degrees. Values greater than 90 degrees indicated that the trajectory had reversed direction between visits to the same regime.

### Evaluating Technical Assumptions

Our homeostatic model makes several simplifying assumptions. First, all channels within a given regulatory mode share the same time constant, and post-translational modifications are removed at the same rate as they are added. In reality, individual channels likely differ in their modification and turnover kinetics, and addition/removal rates may be asymmetric. In fact, the rates of their modification may depend on how the ion channels colocalize with each other (47). However, as long as post-translational changes remain collectively much faster than expression-level changes, the separation of timescales that drives our results should hold. Second, post-translational regulation is implemented solely as midpoint shifts in (in)activation curves with fixed slope, and these shifts are bounded by uniform limits across channels. While biological changes can also alter slope, and the permissible range may differ between channels, these details would alter the magnitude but not the existence of the fast–slow division of labor we observe. Third, we assume that all activity-dependent changes are driven directly by calcium influx, with both regulatory modes sharing the same target activity. In practice, calcium signals can engage distinct downstream pathways, and additional co-regulation or neuromodulatory effects may introduce separate set points for different channels. Such extensions could change the quantitative trajectory of adaptation but would not remove the qualitative role of the faster process in rapid tuning and the slower process in storing persistent adaptations, provided that their characteristic timescales remain well separated.

## DATA AVAILABILITY

The code required to reproduce run the homeostatic model are accessible at github and at Zenodo. Repository: [https://github.com/YugarshiM/fastTimescaleHomeostasis] [DOI: 10.5281/zenodo.12585617]

## ACKNOWLEDGMENTS

The authors acknowledge the support of National Institute of Mental Health (R01MH046742), National Institute of Neurological Disorders and Stroke (R35NS097343), and The Swartz Center for Theoretical Neuroscience at Brandeis University. YM acknowledges valuable discussions with lab members.

